# Maintenance of long-term transposable element activity through regulation by nonautonomous elements

**DOI:** 10.1101/2024.07.13.603364

**Authors:** Adekanmi Daniel Omole, Peter Czuppon

## Abstract

Transposable elements are DNA sequences that can move and replicate within genomes. Broadly, there are two types: autonomous elements, which encode the necessary enzymes for transposition, and nonautonomous elements which rely on the enzymes produced by autonomous elements for their transposition. Nonautonomous elements have been proposed to regulate the numbers of transposable elements, which is a possible explanation for the persistence of transposition activity over long evolutionary times. However, previous modeling studies indicate that interactions between autonomous and nonautonomous elements usually result in the extinction of one type. Here, we study a stochastic model that allows for the stable coexistence of autonomous and nonautonomous retrotransposons. We determine the conditions for this coexistence and derive an analytical expression for the stationary distribution of their copy numbers, showing that nonautonomous elements regulate stochastic fluctuations and the number of autonomous elements in stationarity. We find that the stationary variances of each element can be expressed as a function of the average copy numbers and their covariance, enabling data comparison and model validation. These results suggest that continued transposition activity of transposable elements, regulated by nonautonomous elements, is a possible evolutionary outcome that could for example explain the long co-evolutionary history of autonomous *LINE1* and nonautonomous *Alu* element transposition in the human ancestry.

## 1 Introduction

Transposable elements (TEs) are mobile genetic elements that relocate and replicate in genomes of species across the entire tree of life, from bacteria to animals and plants (Bennett, 2008; Hancks and Kazazian, 2012; Bennetzen and Wang, 2014). They constitute a substantial fraction of eukaryotic genomes, representing 45% of the human genome, over half of the genome in maize and zebrafish, and about one-third in *Drosophila melanogaster* and *Caenorhabditis elegans* (Cosby et al., 2019; Hayward and Gilbert, 2022). Far from being mere remnants of evolutionary history, TEs due to their ability to move, actively shape genomes and may even serve as catalysts of genetic innovations, contributing to the diversification of genes and genomes (Kazazian, 2004).

Nevertheless, TEs are regarded as genomic parasites because they pose potential threats to genomic integrity (Doolittle and Sapienza, 1980) and are sometimes labeled “selfish” due to their proliferation, which may not benefit the host organism and can be deleterious, potentially leading to genomic instability (Orgel and Crick, 1980; Bhat et al., 2022). This instability can arise from: (i) insertions that may disrupt promoters and enhancers, (ii) ectopic recombination between interspersed sequences that could result in deleterious chromosomal rearrangements, (iii) TE insertions into coding regions disrupting their function, or (iv) deleterious effects from TE-encoded complexes or products such as RNA or proteins (Montgomery et al., 1991; Cooke et al., 2014; Tubio et al., 2014). Based on these characteristics, a TE life-cycle in genomes has been proposed (Le Rouzic et al., 2007): after TE invasion and establishment, TEs proliferate in the host genome until they become silenced or domesticated by the host. In each outcome, domestication or silencing, TEs lose their transposition ability. Yet, examples of TEs exist that have maintained their transposition ability over millions of years. One of the most prominent examples are *LINE1* and *Alu* elements in the human genome lineage (Kazazian, 2004).

Three different explanations for such continued TE activity have been proposed (Brookfield and Badge, 1997). The first mechanism is referred to as transposition-selection balance (e.g. Charlesworth and Charlesworth, 1983; Charlesworth, 1991). Due to the negative fitness effects of TEs on their host, individuals with large TE copy numbers are selectively disadvantageous and produce less offspring than hosts with fewer numbers of TEs. Therefore, transposition of TEs in the genome is balanced by selection against large TE copy numbers on the host level, thus resulting in stable TE copy numbers over time. The second regulation mechanism is TE self-regulation (e.g. Charlesworth and Charlesworth, 1983; Charlesworth and Langley, 1986). Here, the transposition rate of TEs decreases with TE copy number. This is reminiscent of ecological population models like logistic growth, the transposition rate corresponding to the growth rate and the TE copy number to the population size. This TE self-regulation will also result in stable copy numbers over time. The last regulation mechanism that has been suggested is based on an interaction between *autonomous* and *nonautonomous* TEs.

Autonomous TEs encode the entire transposition machinery necessary for their transposition. In contrast, nonautonomous TEs rely on the transposition machinery of autonomous elements, which is why they have been characterized in the literature as “hyperparasites” (Robillard et al., 2016), “superparasites” (Startek et al., 2013), and “parasites of parasites” (Suh, 2019). Examples of autonomous-nonautonomous TE pairs include the already mentioned *LINE1* and *Alu* elements in the human genome (Boeke, 1997; Dewannieux et al., 2003; Wagstaff et al., 2012), RLG *Wilma* and RLG *Sabrina* in the wheat genome (Wicker et al., 2021), activator (Ac) and dissociation (Ds) elements in the maize genome (McClintock, 1956), or Tc1/mariner and *Stowaway* Miniature Inverted Repeat Transposable Elements (MITEs) in the rice genome (Feschotte et al., 2003). TEs are additionally characterized based on their transposition mechanisms into retrotransposons, which use a copy-and-paste mechanism, and DNA transposons, which use a cut-and-paste mechanism. Retrotransposons are further categorized by their chromosomal integration mechanisms into LTR, TEs with long terminal repeats, and non-LTR retrotransposons. In this manuscript, we will model non-LTR retrotransposons.

Various theoretical models have been developed to investigate the interactions between autonomous and nonautonomous retrotransposons. Interestingly, essentially all of these studies find that coexistence of nonautonomous and autonomous TEs is typically not possible over long periods of time (e.g. Brookfield, 1996; Le Rouzic and Capy, 2006; Startek et al., 2013), though some deterministic and stochastic simulation examples show that long-term coexistence is possible (Le Rouzic and Capy, 2006). Many of these studies model a nonautonomous TE as arising through mutations from the autonomous TE, making the two TE types related to each other, in the sense of sequence similarity. Moreover, all of these models assume deleterious fitness effects of TEs on the host. But even if the nonautonomous TE is ‘unrelated’ to the autonomous TE, no stable coexistence was found in simulations. A theoretical explanation and exploration of whether and when continued transposition of an autonomous-nonautonomous TE pair is possible, as observed for the seemingly unrelated *LINE1* and *Alu* elements, is therefore missing.

Here, we investigate this discrepancy between theoretical prediction and empirical observation. One detail that has not been considered explicitly in previous studies is the interaction between the two TE types on the molecular level. To account for this, we explicitly model the molecular processes of transposition and study two ‘unrelated’ non-LTR TE types, following the model by Xue and Goldenfeld (2016). In this model, TEs do not affect host fitness and the dynamics on the host level are neutral. We study the stability of and stochastic fluctuations around the TE coexistence equilibrium. Importantly, our theoretical analysis provides a parameter-independent prediction about TE copy number variation in a host genome, making it suitable for comparison with empirical data. Lastly, we investigate the effect of recombination on TE copy numbers and compare our prediction to stochastic simulations. Theory and simulations agree for low recombination rates, but we underestimate the variance between the TE types for large recombination rates. This means that our theory is still useful for data comparison with TEs that accumulate in genomic regions with suppressed recombinations, examples of which are abundantly described in the literature (e.g. Dolgin and Charlesworth, 2008; Errbii et al., 2024).

We start by describing the model and analyzing the coexistence dynamics of the two TE types in a single lineage, which translates to the population distribution of TE copy numbers. Second, we explore how random mating and recombination affects TE copy numbers in a host population.

## 2 Model

We study the dynamics of an autonomous-nonautonomous non-LTR TE pair within a genome. We first describe the processes that are underlying the changes in TE copy number and derive a pair of stochastic differential equations that model these dynamics. We then extend the model to a host population to study the effects of random mating and recombination on the TE dynamics. This requires a more detailed description of the host’s genetic setup.

### 2.1 TE dynamics

The current mechanistic understanding of non-LTR autonomous and nonautonomous retrotransposon mobilization can be summarized as follows (Dewannieux et al., 2003). Both autonomous and nonautonomous TEs are transcribed, but only autonomous transcripts encode the proteins necessary for retrotransposition. When an autonomous transcript is translated into protein in the ribosome, that very transcript tends to bind to the protein, likely through recognition of its poly-A tail. This binding can result in retrotranscription into another genomic site, a phenomenon known as *cis* replication (Wei et al., 2001). Alternatively, other transcripts can also bind to the nascent protein, a process referred to as *trans* replication. Importantly, a nonautonomous transcript co-localizing at the same ribosome can bind to the nascent protein due to sequence similarity in its poly-A tail with the autonomous transcript, which it then uses for retrotranscription elsewhere in the genome (Boeke, 1997; Dewannieux et al., 2003).

To translate the biology into mathematical formulas, we denote the copy number of active autonomous and nonautonomous TEs within a genome at a given time *t* as *X*(*t*) and *Y* (*t*), respectively. Let *C* represent the complex of autonomous TE-encoded mRNA and nascent protein. An autonomous TE, *X*, encodes this complex which forms at a rate *a*. Subsequently, the complex retrotransposes producing a new *X* element at a rate *b*_1_ (*cis* replication). Moreover, the complex *C* can be hijacked by a nonautonomous TE, *Y*, to replicate itself at a rate *b*_2_ (*trans* replication). Lastly, the complex fails to produce a new autonomous or nonautonomous TE at a rate *d*_2_. The number of (active) TEs decreases either by deactivation, e.g. through host silencing or mutations, or excision from the genome. We lump these degradation processes in the parameter *d*_1_. TE-type specific degradation rates are investigated in Appendix D. These dynamics are illustrated in Fig. 1 and yield the following model, previously described by Xue and Goldenfeld (2016):

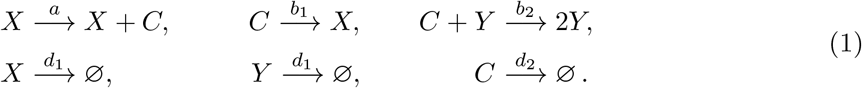

**Figure 1.**
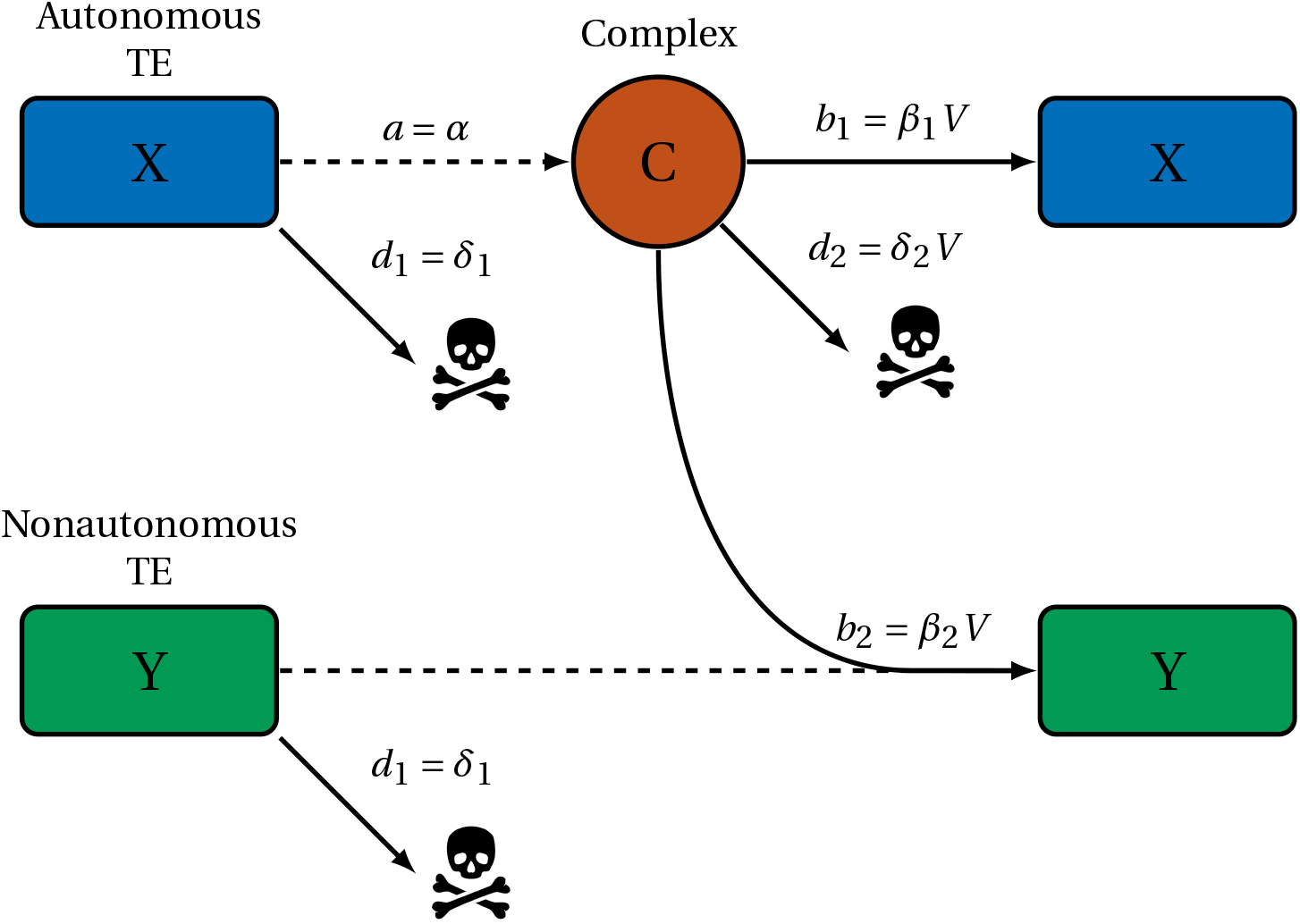
Schematic of autonomous-nonautonomous TE interaction. The diagrammatic description illustrates the interaction between autonomous and nonautonomous TEs. The autonomous TE, *X*, encodes the complex, *C* at a rate *a* = *α*, and this complex retrotranscribes to produce a new *X* element at rate *b*_1_ = *β*_1_*V* . The nonautonomous TE, *Y*, hijacks the complex produced by the autonomous TE to duplicate itself at the rate *b*_2_ = *β*_2_*V* . Both types of TEs deactivate at rate *d*_1_ = *δ*_1_, and the complex fails to produce a new TE (essentially a degradation process) at rate *d*_2_ = *δ*_2_*V* . Solid arrows indicate transformations that involve a change in numbers at the start and end of the arrow, and dashed arrows indicate production, where only the numbers at the tip of the arrow change.

TE numbers in the germline change slowly from generation to generation in a host species population (Suh et al., 1995). Compared to this evolutionary time scale, the formation and degradation of the intermediate complex occur on a much faster time scale. Alternatively, TE numbers can also change in the somatic cells, where the same distinction between fast complex and slow TE processes is appropriate. Mathematically, this translates to dynamics on two different time scales, which can be separated. To achieve this separation, we scale the number of TEs by the parameter *V*, which represents the order of magnitude of the number of TEs (or the length of the genome that the TEs occupy). The number of complexes is of order 1 ≪*V* . We then study the rescaled variables *x*^*V*^ = *X/V* and *y*^*V*^ = *Y/V* while the number of complexes remains on the original scale. Following classical mass action kinetics, we also transform the binary reaction rate *b*_2_ by multiplying it with 1*/V*, which now describes the probability of interaction between a complex and a nonautonomous TE; for a more detailed discussion of this classical scaling, we refer to Section 3.1 in Anderson and Kurtz (2011). To account for the relatively faster dynamics of the complex, we scale the rates as follows: *a* = *α, b*_1_ = *β*_1_*V*, *b*_2_ = *β*_2_*V*, *d*_1_ = *δ*_1_, *d*_2_ = *δ*_2_*V* . The scaling indicates that the formation of the complex at rate *a* and deactivation of the TEs at rate *d*_1_ occur on the same (slow) time scale. Additionally, the scaling shows that once the complex is formed, the production of new autonomous and nonautonomous TEs at rates *b*_1_ and *b*_2_, respectively, occur on the same (fast) time scale as the rate *d*_2_ at which no TE is retrotranscribed by the complex.

The stochastic dynamics of this rescaled system are then given by:

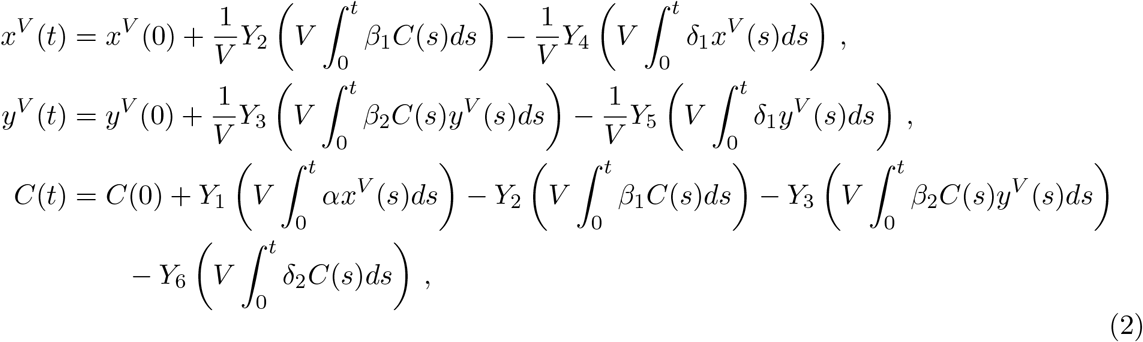

where *Y*_*i*_, *i* ∈ {1, …, 6} are independent rate-1 Poisson processes corresponding to the reactions in Eq. (1).

For *V* sufficiently large, this system exhibits a time-scale separation, where the complex dynamics, *C*(*t*), are occurring much faster than the dynamics of the two TE types, *x*^*V*^ and *y*^*V*^ . Consequently, given a certain TE configuration, the complex reaches a quasi-steady state (Ball et al., 2006). This equilibrium of the complex is given by

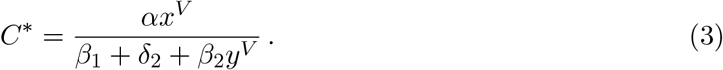

To derive the stochastic differential equations (SDEs) corresponding to Eq. (2), we employ the infinitesimal generator and convergence results of density dependent Markov processes (Ethier and Kurtz, 1986, Chapters 4 & 11). We find

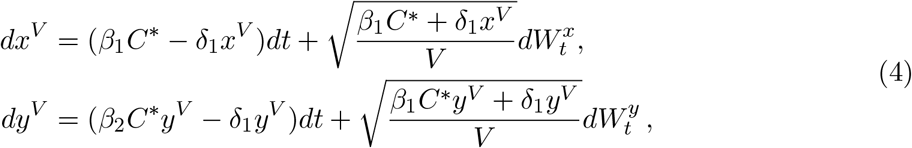

where 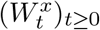 and 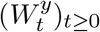 are two independent standard Brownian motions, and the stochastic differential is interpreted in the sense of It ô. Inserting the equilibrium value of the complex gives

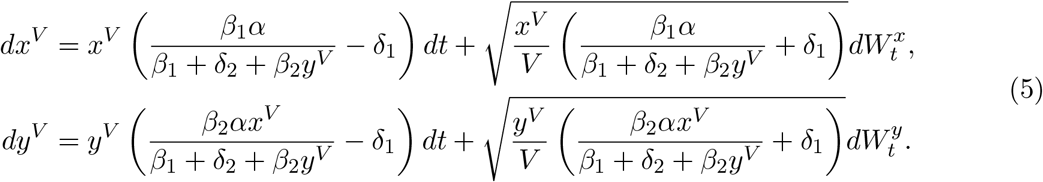

This system of stochastic differential equations is the basis of the subsequent analysis to derive our results.

### 2.2 Model of TE dynamics in a population

We now extend the model to capture the TE copy number dynamics in a host population. We model a diploid population of fixed size *N* with non-overlapping generations (Wright-Fisher model). Each individual has a single pair of chromosomes. Each chromosome contains 5000 potential integration sites for TEs. The population is initialized with 200 autonomous and 200 nonautonomous randomly placed TEs per chromosome in each individual.

For each individual in the next generation, two-parent individuals are randomly selected from the population. Our model assumes that TEs impose no fitness cost, meaning all individuals have equal chances of reproducing, and thus the dynamics are neutral. A haploid gamete is generated from the genome of each parent. In the case of recombination, the number of crossovers per chromosome is drawn for different cases from a Poisson distribution whose means are a total map distance of 1, 10, and 100 cM between extreme loci. These total map distances correspond to recombination rates of 0.01, 0.1 and close to 0.5 respectively (Figure 2 in Pealba and Wolf, 2020). The locations of crossover events are uniformly distributed along the chromosomes. From each pair of recombinant chromosomes, one is randomly selected to form the gamete. During this stage, transposition and deletion events for both autonomous and nonautonomous elements occur, i.e., TE dynamics happen during the haploid stage of the life cycle. The numbers of transposition and deletion events follow Poisson distributions, with the means determined by the transposition and deletion rates, the corresponding stationary number of complexes, and the current number of TEs in the genome. Once both parents have produced gametes and TE dynamics occurred, the gametes of the two parent individuals are combined to form a new diploid individual.

**Figure 2.**
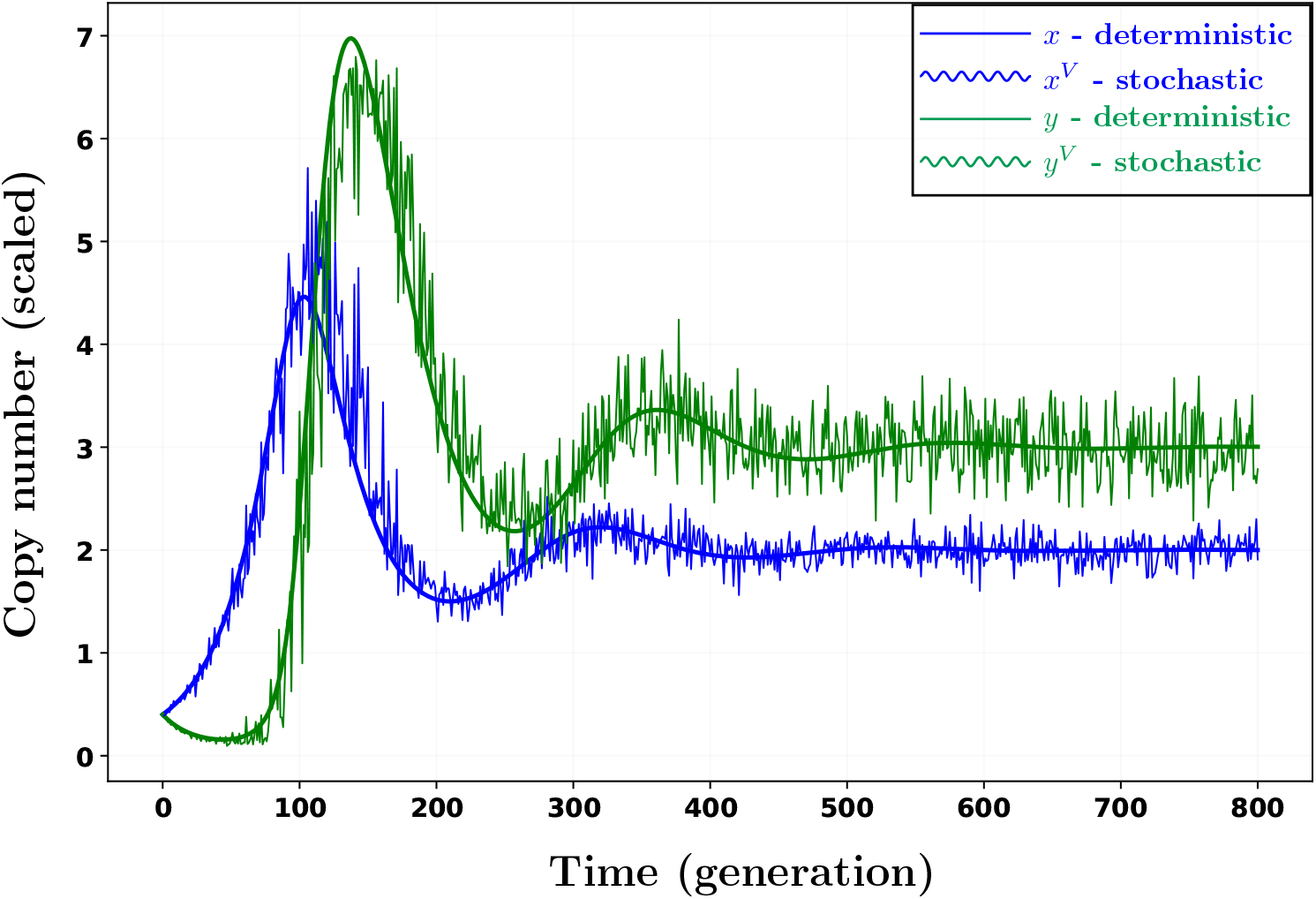
Simulation of the autonomous-nonautonomous TE interaction. The graph shows the deterministic (straight lines) and stochastic (wavy lines) dynamics of active autonomous (blue) and nonautonomous (green) TE copy numbers over time. These dynamics are shown for a single chromosome in a population of 1000 diploid individuals. The TEs cyclically approach their equilibrium and then fluctuate around their respective equilibrium. Parameter values are *α* = 2 × 10^−1^, *β*_1_ = 1 × 10^−2^, *β*_2_ = 5 × 10^−3^, *δ*_1_ = 5 × 10^−2^, *δ*_2_ = 1.5 × 10^−2^, *r* = 0, and *V* = 500.

### 2.3 Stochastic simulations

To update TE numbers in the haploid stage of the life cycle, we implemented the TE dynamics model (Eq. 1) using the Euler tau-leaping method (Gillespie, 2001). For comparison, we also simulated the corresponding deterministic model for each TE type (deterministic part in Eq. (5)), see Fig. 2. All simulations were executed in Julia, while all subsequent data analyses and plotting were performed in Python. The source codes and data files to reproduce the figures and simulation results are available at https://github.com/Ade-omole/TEs_long_term. Our default parameter set is stated in Table 1. We note that these parameters are higher than what would be expected empirically. For example, the transposition rate and TE deletion rate are estimated to be of order 10^−3^ − 10^−5^ (Suh et al., 1995). However, this deviation from empirical evidence does not affect our general, parameter-independent results and speeds up simulations.

**Table 1:**
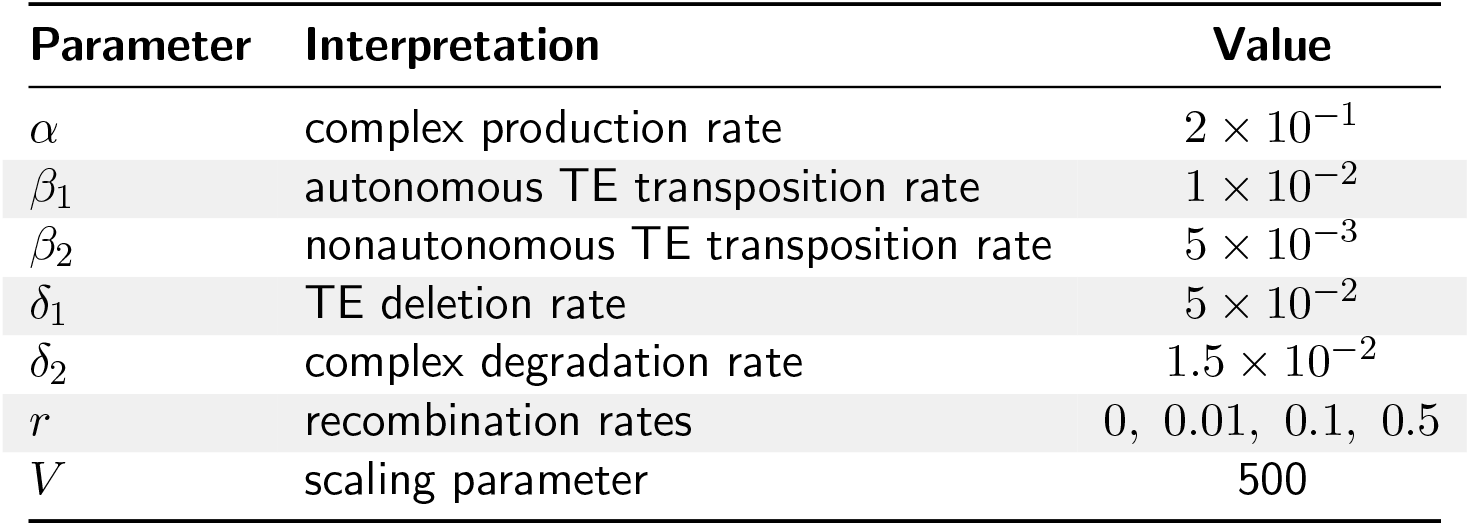
Overview of parameters. We summarize all parameter abbreviations, their biological interpretations and default values in the simulations.

## 3 Results

We now analyze the system of stochastic differential equations in Eq. (5) and numerically assess the validity and robustness of the theoretical analyses. We start by investigating the deterministic model, which exhibits a locally stable coexistence state with both autonomous and nonautonomous TEs under certain parameter configurations. We then study the stochastic dynamics and describe the stationary distribution by computing the fluctuations around the deterministic equilibrium. Lastly, we compare the theoretical prediction to stochastic simulations and study the effect of recombination on TE copy numbers.

### 3.1 Stable coexistence of TE types

#### Deterministic equilibrium

The deterministic dynamics of the number of TEs from Eq. (5) are obtained by letting *V* → ∞. Writing 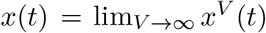 and 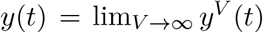, we obtain

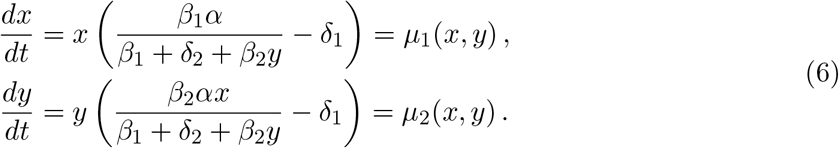

Setting both derivatives equal to zero, the equilibrium of system (6) is found as

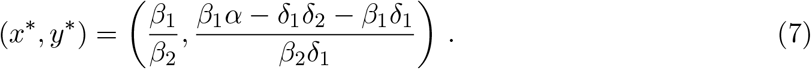

A non-trivial fixed point of the system, wherein both TE numbers are positive, exists if the parameters satisfy the following condition:

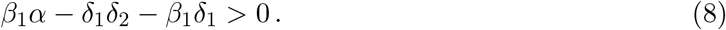

#### Local stability analysis

To analyze the local stability of the equilibrium described in Eq. (7), we compute the Jacobian matrix of the dynamics outlined in Eq. (6) at the equilibrium, given by

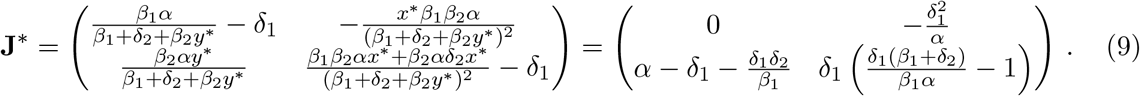

Using the determinant-trace condition (e.g. Otto and Day, 2007, page 238), the eigenvalues of **J**^*^ have negative real part if det(**J**^*^) > 0 and tr(**J**^*^) < 0. Both conditions are satisfied if and only if the equilibrium is feasible, which is the case if Eq. (8) holds. Thus, the non-trivial equilibrium is locally stable if it exists.

To gain intuition about the condition for feasibility and local stability of the coexistence equilibrium, we rewrite condition (8) as follows:

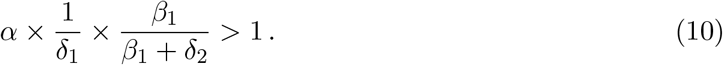

The second term (1*/δ*_1_) is the average life-time of an autonomous TE. Therefore, the first product *α/δ*_1_ reflects the average number of produced complexes by a single autonomous TE until it is excised or becomes inactive. The last term (ratio *β*_1_*/*(*β*_1_ + *δ*_2_)) is the probability that a complex produces an autonomous TE in the absence of nonautonomous TEs. The left-hand side of Eq. (10) can therefore be identified as the reproduction number of autonomous TEs in the absence of nonautonomous TEs. Interestingly, this implies that the condition for coexistence of autonomous and nonautonomous TEs depends solely on the parameters of the autonomous TEs, but note that the autonomous TE copy number depends on the nonautonomous transposition rate. This is because condition (10) simplifies to *β*_1_*α/*(*β*_1_ + *δ*_2_) − *δ*_1_ > 0, which is *µ*_1_(*x*, 0) in Eq. (6). Therefore, coexistence of the two TEs in this model is possible and (locally) stable if autonomous TEs grow exponentially in the absence of nonautonomous TEs. We note that this exponential growth would in reality eventually be stopped either by host regulation or deleterious fitness effects due to TE insertions, both of which we do not consider in our model (Kelleher et al., 2020; Charlesworth and Charlesworth, 1983)

The nonautonomous TE type plays a regulatory role in this interaction of the two TEs. This is apparent in Eq. (7), where *β*_2_ scales the equilibrium TE copy number of the autonomous TE: the larger the rate of nonautonomous TE production, *β*_2_, the smaller is the equilibrium value of the autonomous TE, *x*^*^. This implies that although the nonautonomous TE does not directly contribute to the stable coexistence of the two TEs through its own transposition, it plays a pivotal role in regulating TE proliferation.

#### Relative copy numbers of the TE types

The copy numbers of the two TE types rely on the average number of complexes produced by the autonomous TEs. The conditions governing the relative copy numbers of the autonomous and nonautonomous TEs derived from Eq. (7) are as follows:

##### Proposition 1

*If the equilibrium is feasible (and therefore locally stable), then*

a. *if* 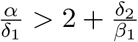,*then x* ^*^ < *y*^*^,
b. if 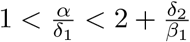,*then x* ^*^ < *y* ^*^,
c. *if* 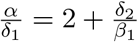,*then x* ^*^ = *y*^*^,

The proof is stated in Appendix A.

The proposition provides additional insights into the relationship between the two types of TEs: The autonomous TE is more or less abundant than the nonautonomous TE depending on whether the average number of complexes produced by an autonomous TE before its deactivation, *α/δ*_1_, is smaller or larger than the term 2 + *δ*_2_*/β*_1_. This term depends on the transposition rate of the autonomous TE but is independent of the nonautonomous transposition rate. If the transposition rate of the autonomous TE, *β*_1_, is low, the term 2 + *δ*_2_*/β*_1_ becomes large. This means that the average number of complexes required for the autonomous TE to be less abundant than the nonautonomous TE, as in (a), increases. Conversely, if the transposition rate of the autonomous TE, *β*_1_, is high, the term 2 + *δ*_2_*/β*_1_ approaches the value 2. This implies that the average number of complexes required for the autonomous TE to be less abundant is low, but above 2. This means that the parameter space for autonomous TEs to be less abundant than nonautonomous TEs decreases with decreasing autonomous transposition rate *β*_1_, Fig. 6 in Appendix A. At the same time, the inverse conclusion holds true for the condition for autonomous TEs to be more abundant than nonautonomous TEs (part (b) of Proposition 1).

### 3.2 Stationary distribution of the TEs

We now examine the stochastic fluctuations of the stochastic process around the deterministic equilibrium. There are different approaches to study these fluctuations in systems with time-scale separations. The most accurate method involves deriving a central limit theorem for the stochastic process defined in Eq. (2) (Kang et al., 2014). Alternatively, a simpler approach is based on the Langevin equation (van Kampen, 2007), which uses the quasi-steady state assumption at every step, i.e., only the stationary number of complexes is taken into account. Here in the main text, we use the Langevin equation approach, also called the linear noise approximation. The analogous analysis using the central limit theorem is relegated to Appendix B and yields almost the same result. The linear noise approximation derives the distribution of the process from the local dynamics around the deterministic equilibrium points through a Taylor expansion; we follow the exposition from Czuppon and Traulsen (2021). To this end, we study the fluctuations of two TE types around their respective equilibrium. The stochastic processes characterizing these fluctuations, denoted by *U*_1_ and *U*_2_, are defined as:

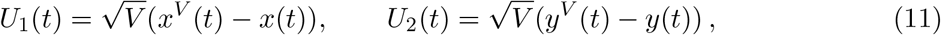

where *x*(*t*) and *y*(*t*) represent the deterministic processes (Eq. (6)) associated to the stochastic processes *x*^*V*^ (*t*) and *y*^*V*^ (*t*) (Eq. (5)). Denoting **U** = (*U*_1_, *U*_2_)^⊤^, the dynamics of the stochastic fluctuations can be approximated by

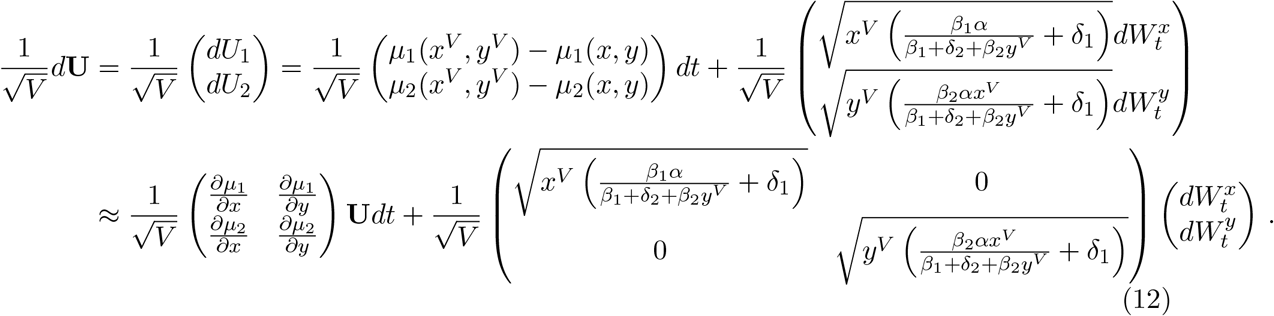

Inserting the deterministic coexistence equilibrium (Eq. (7)), we obtain

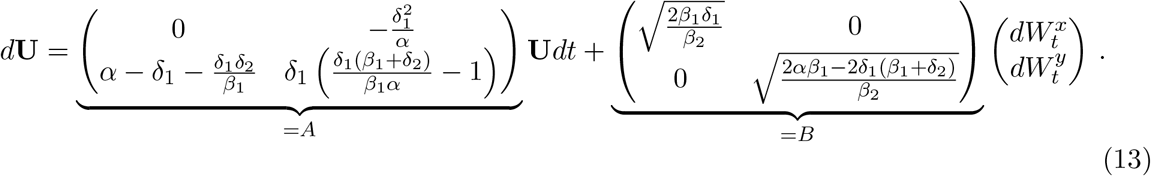

This stochastic process is a two-dimensional Ornstein-Uhlenbeck process. Its stationary covariance matrix Σ reads (Czuppon and Pfaffelhuber, 2018, Theorem 3)

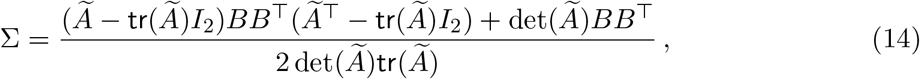

where Ã = −*A* and *I*_2_ is the two-dimensional identity matrix. Inserting the matrices as defined in Eq. (13) we find the rescaled stationary covariance matrix 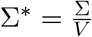

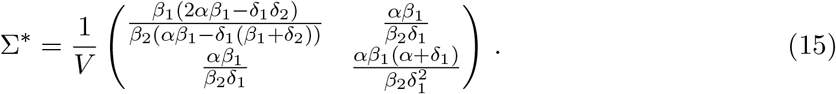

Writing *ξ* = *αβ*_1_*/*(*β*_2_*δ*_1_) for the covariance term, we can rewrite this as

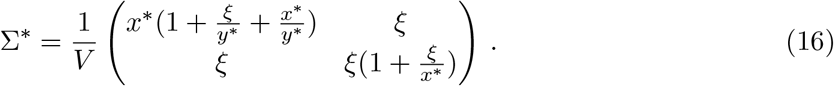

Plugging everything into Eq. (13) and rearranging, we find that the stationary distribution of the two-dimensional TE numbers, which we denote by *ψ*^*^, is given by

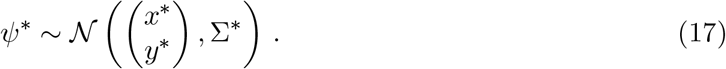

The stationary variance of the autonomous TE, as indicated by the covariance matrix in Eq. (16), depends on its own mean, *x*^*^, the mean of the nonautonomous TE, *y*^*^, and the covariance of the TEs in stationarity, *ξ*. The stationary variance of the nonautonomous TE, however, depends only on the covariance, *ξ*, and the mean value of the autonomous TE, *x*^*^, but not on its own mean, *y*^*^. Additionally, we note that it substantially depends on the covariance. The covariance itself, given by *αβ*_1_*/β*_2_*δ*_1_ = *αx*^*^*/δ*_1_, can be interpreted as the overall average number of complexes produced in stationarity. This implies that the stationary variance of nonautonomous TEs is strongly affected by the complex (or transposition machinery) encoded by the autonomous TE.

Note that in Eq. (17) we have computed the stationary distribution along a single trajectory of TE dynamics, i.e., we have computed the temporal fluctuations of TE copy numbers. These fluctuations are the same as the distribution of TE copy numbers in a population at a single point in time, at least approximately for large population sizes. This is a mathematical property (ergodicity) of the Ornstein-Uhlenbeck process, which allows to translate statistical properties between temporal (TE dynamics along a single lineage) and spatial (host population) scales (Chapter 8 in Kallenberg, 2021).

In Fig. 3 we compare our theoretical prediction of the stationary distribution to stochastic simulations and find a very good fit. We also plot the stationary distribution found through the central limit theorem (Eq. (24) in Appendix B) and see that for our parameters there is no visible difference between the two distributions. In fact, we find that the stationary variance of the nonautonomous TE and covariance are exactly the same. The stationary variance of the autonomous TE differs on the scale *O*(1*/V* ) (Eq. (25) in Appendix B). In Fig. 3, we also compute an empirical stationary distribution, where we estimate the equilibrium TE values, *x*^*^ and *y*^*^, and the covariance *ξ* from the data and then use Eq. (16) to compute the stationary variances. The empirical stationary distribution fits well with the analytical prediction and data for both TE types.

**Figure 3.**
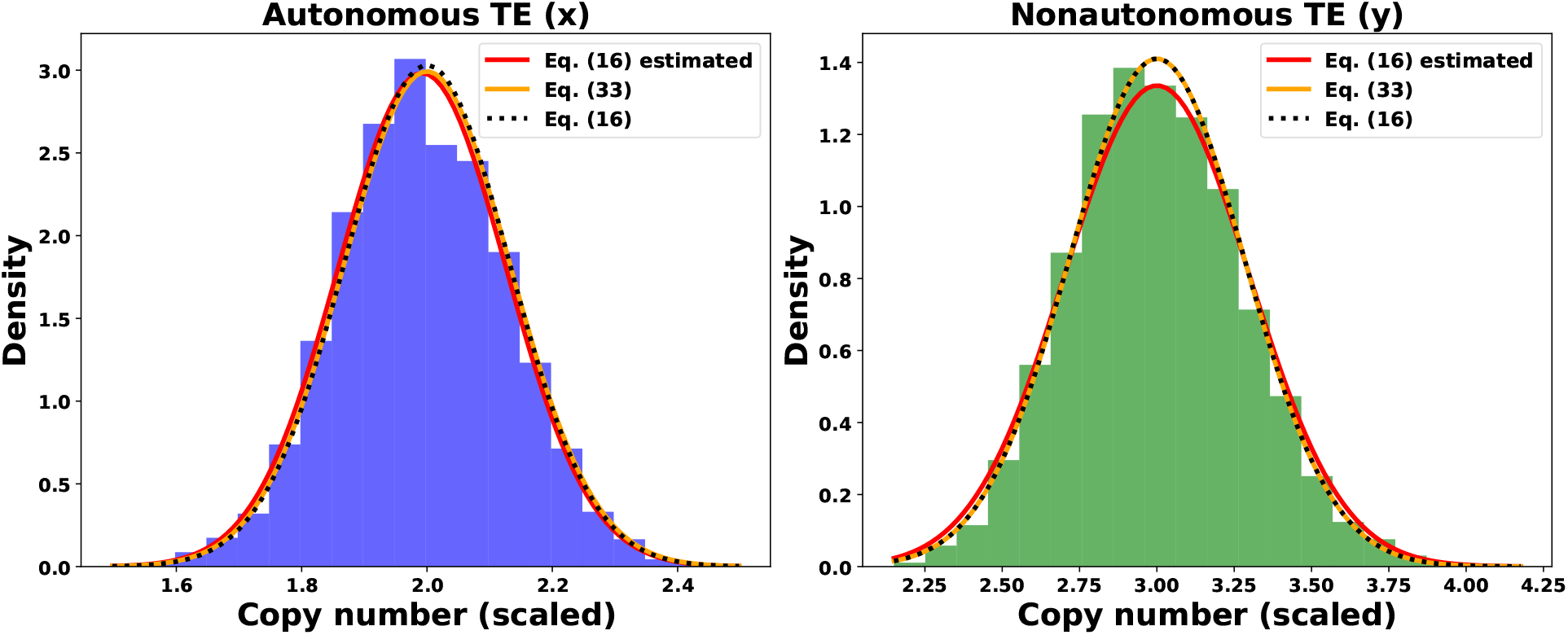
Comparison of theoretical stationary distributions of TE numbers to stochastic simulations under no recombination. Autonomous (left) and nonautonomous (right) TE number histograms are obtained from the generation 1000 of a single stochastic simulation run of population size 10^4^, with no recombination (the equilibrium is already reached at that time, see Fig. 2). Parameter values are the standard parameter set (Table 1), resulting in theoretical mean TE copy numbers of *x*^*^ = 2 and *y*^*^ = 3 (Eq. (7)) and theoretical TE stationary variances of 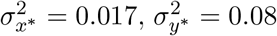 (Eq. 16) and 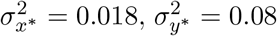 (Eq. 24). The dashed, black line shows the linear noise approximation from the main text, and the stationary variance computed with the central limit theorem is shown by solid orange lines. Additionally, we plot the estimated stationary distribution (red, solid) by estimating the means 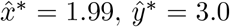, and covariance 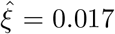 from the data, then computed with Eq. (16) the estimated variances 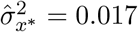 and 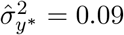. For comparison, the theoretical covariance is *ξ* = 0.016.

Similar to the comparison of mean values of TEs in stationarity, we now provide analogous statements for the stationary variances. We denote the stationary variances of the autonomous and nonautonomous TEs as 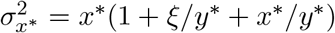 and 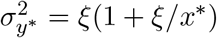, respectively.

#### Proposition 2

*Let the coexistence equilibrium* (*x*^*^, *y*^*^) *be locally stable. Then we have:*

a. *If x*^*^ ≤ *y*^*^, *then* 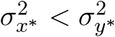.
b. *Set* 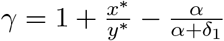 *and let x*^*^ > *y*^*^. *We then have:*

(i) *If* 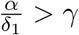,*then* 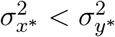,
(ii) *If* 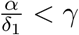,*then* 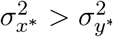,
(iii) *If* 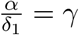,*then* 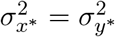.

The proof is provided in Appendix C.

Proposition 2 shows the relationship between the stationary variances of the two types of TEs and their respective mean values. Importantly, we find that the stationary variance of the nonautonomous TE is always larger than that of the autonomous TE if the nonautonomous TE has a larger equilibrium value. The case where the mean copy number of the autonomous TE is larger than that of the nonautonomous TE is more involved, i.e., no simple prediction about the relationship of the stationary variances is possible. Fig. 4 visualizes these different scenarios as we vary the complex production rate *α*.

**Figure 4.**
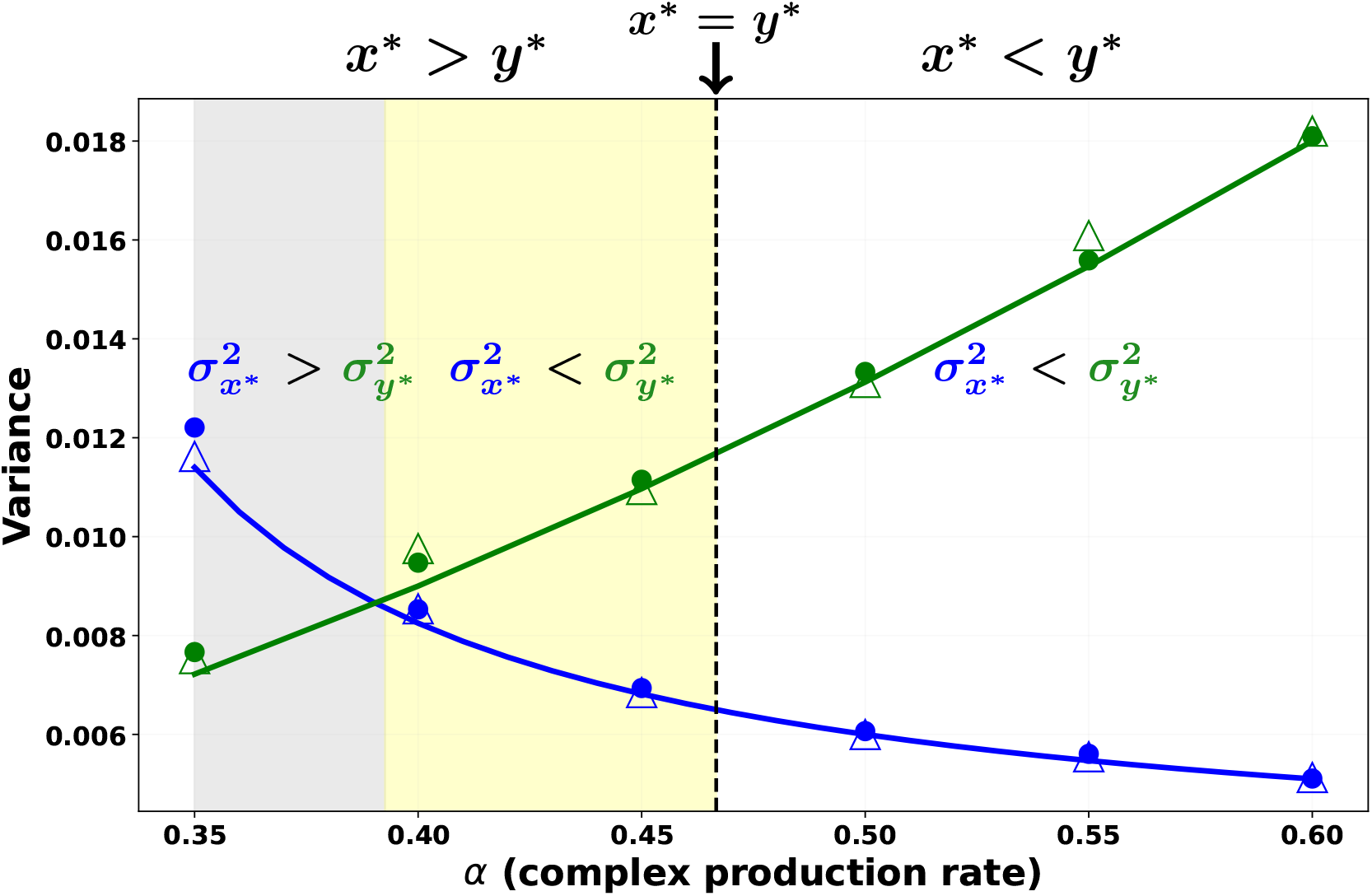
Graphical representation of Proposition 2. The figure shows simulated data (circle), analytic (line) and estimated (triangle) variances of the autonomous (blue) and nonautonomous (green) TEs for constant parameter values *β*_1_ = 0.3, *β*_2_ = 0.4, *δ*_1_ = 0.2, *δ*_2_ = 0.1, and varying alpha values, as indicated. The figure shows that the relative copy numbers of the TEs depend on the rate of complex production by the autonomous TE, *α*. If the mean copy number of the autonomous TE is less than that of the nonautonomous TE (right of the dashed line), then the autonomous TE’s stationary variance is less than that of the nonautonomous TE (white background). Conversely, if the mean copy number of the autonomous TE is greater than that of the nonautonomous TE (left of the dashed line), then the stationary variance of the autonomous TE can be less (yellow), equal, or greater (gray) than that of the nonautonomous TE, depending on the threshold value of *γ* = 0.39 for our parameter set.

### 3.3 Effect of recombination

In this section, we investigate the effect of recombination on the TE dynamics, particularly the stochastic fluctuations around their equilibria. In Fig. 5, we observe that the TE equilibria, or mean copy numbers, are not affected under recombination compared to no recombination (see Fig. 3). A summary of stationary mean and variance values is provided in Table 2. Moreover, to examine the impact of recombination on the variances and covariance of the TE types, we compare the theoretical stationary distribution and the estimated distribution with the data from stochastic simulations under recombination rates *r* = 0.01, *r* = 0.1, and free recombination, i.e. *r* = 0.5 (Fig. 5). Under a low recombination rate (Fig. 5a, *r* = 0.01), the estimated and analytical predictions fit well with the histogram from the simulated data for both TE types. However, as the recombination rate increases up to free recombination (Fig. 5b, c), the estimated stationary distribution for the autonomous TE fits the simulated data, but the analytical prediction shows a continuous loss of fit due to a decrease in data variance. For the nonautonomous TE, there is a progressive loss of fit between both the estimated and analytical stationary distributions and the simulated data as the recombination rate increases. This is attributed to a continuous decrease in the estimated covariance due to recombination, and consequently, the data variance. Importantly, we expect this to depend on the relationship of overall TE activity and the recombination rate. Broadly speaking, if the recombination rate is larger than the TE activity, we expect that recombination breaks TE dependence faster than it accumulates due to transposition events. Thus, high recombination rates substantially reduce the covariance between genomic sites, which results in a loss of fit.

**Table 2:** Comparison of parameter estimates with recombination rates. The table shows the autonomous and nonautonomous TE mean, variance, and covariance estimates from Figs. 3 and 5 under different recombination rates (*r*).

| | $r = 0$ | $r = 0.01$ | $r = 0.1$ | $r = 0.5$ |
| --- | --- | --- | --- | --- |
| $\hat{x}^*$ | 1.99 | 2.00 | 1.99 | 1.99 |
| $\hat{y}^*$ | 3.0 | 3.00 | 2.99 | 3.01 |
| $\hat{\sigma}_{x^*}^2$ | 0.017 | 0.017 | 0.012 | 0.007 |
| $\hat{\sigma}_{y^*}^2$ | 0.09 | 0.07 | 0.026 | 0.0005 |
| $\hat{\xi}$ | 0.017 | 0.015 | 0.008 | 0.0004 |

**Figure 5.**
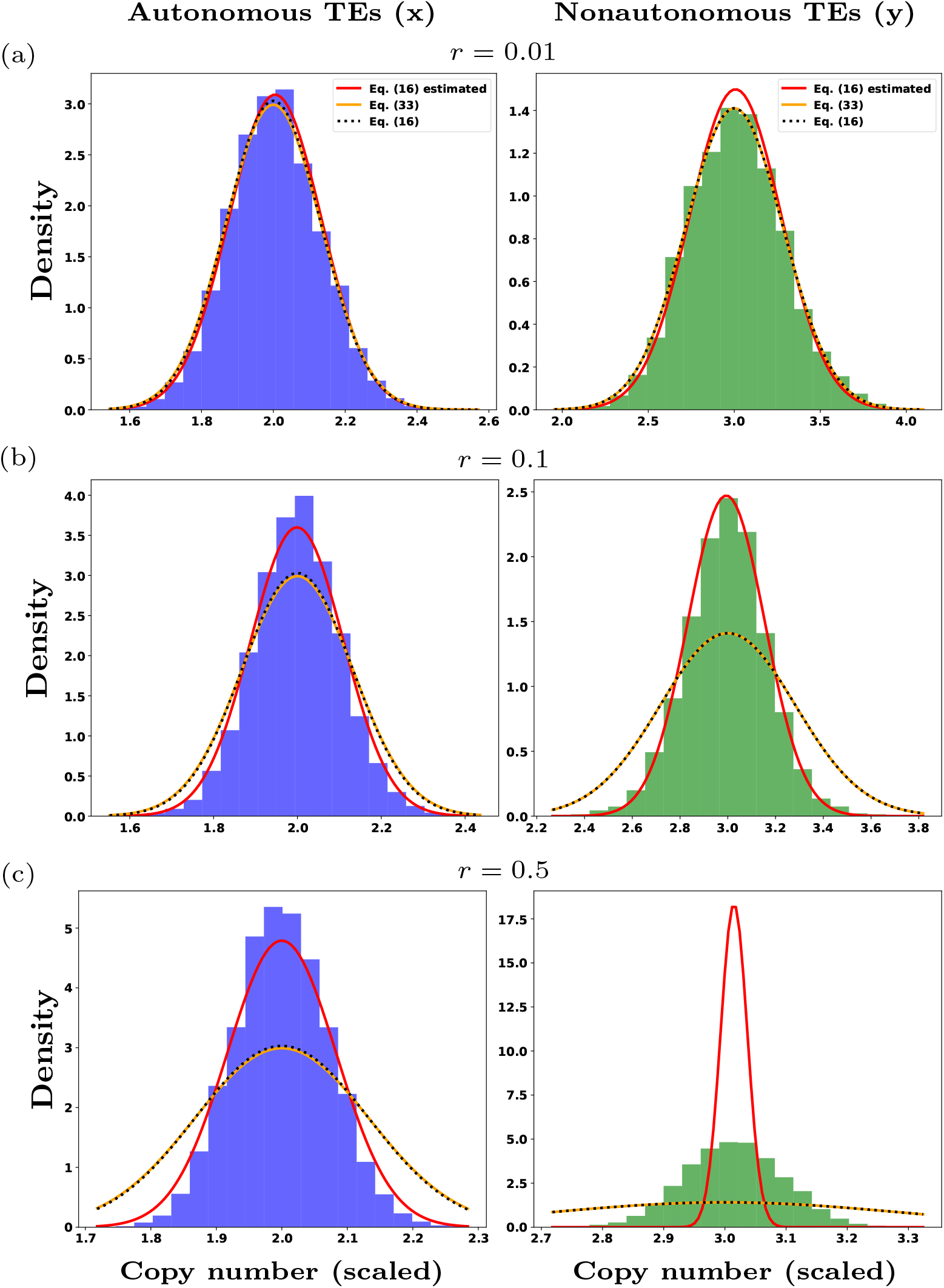
Comparison of theoretical stationary distributions of TE numbers to stochastic simulations under recombination. Autonomous (left) and nonautonomous (right) TE copy number histograms are obtained from generation 1000 of a single stochastic simulation run of population size 10^4^, under recombination rates **(a)** 0.01, **(b)** 0.1, and **(c)** 0.5. Parameter values are the standard parameter set (Table 1), resulting in theoretical mean of *x*^*^ = 2, *y*^*^ = 3 (Eq. (7)), theoretical stationary variances of 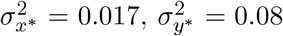 (Eq. (16)) with 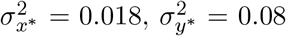 (Eq. (24)), and theoretical covariance of *ξ* = 0.016. The dashed, black line shows the linear noise approximation from the main text, and solid orange lines show the stationary variance computed with the central limit theorem. Additionally, we plot the estimated stationary distribution (red, solid) by estimating the means 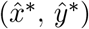, and covariance 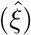 from the data, then computed with Eq. (16) the estimated variances 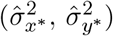 .The estimates are shown in Table 2.

**Figure 6.**
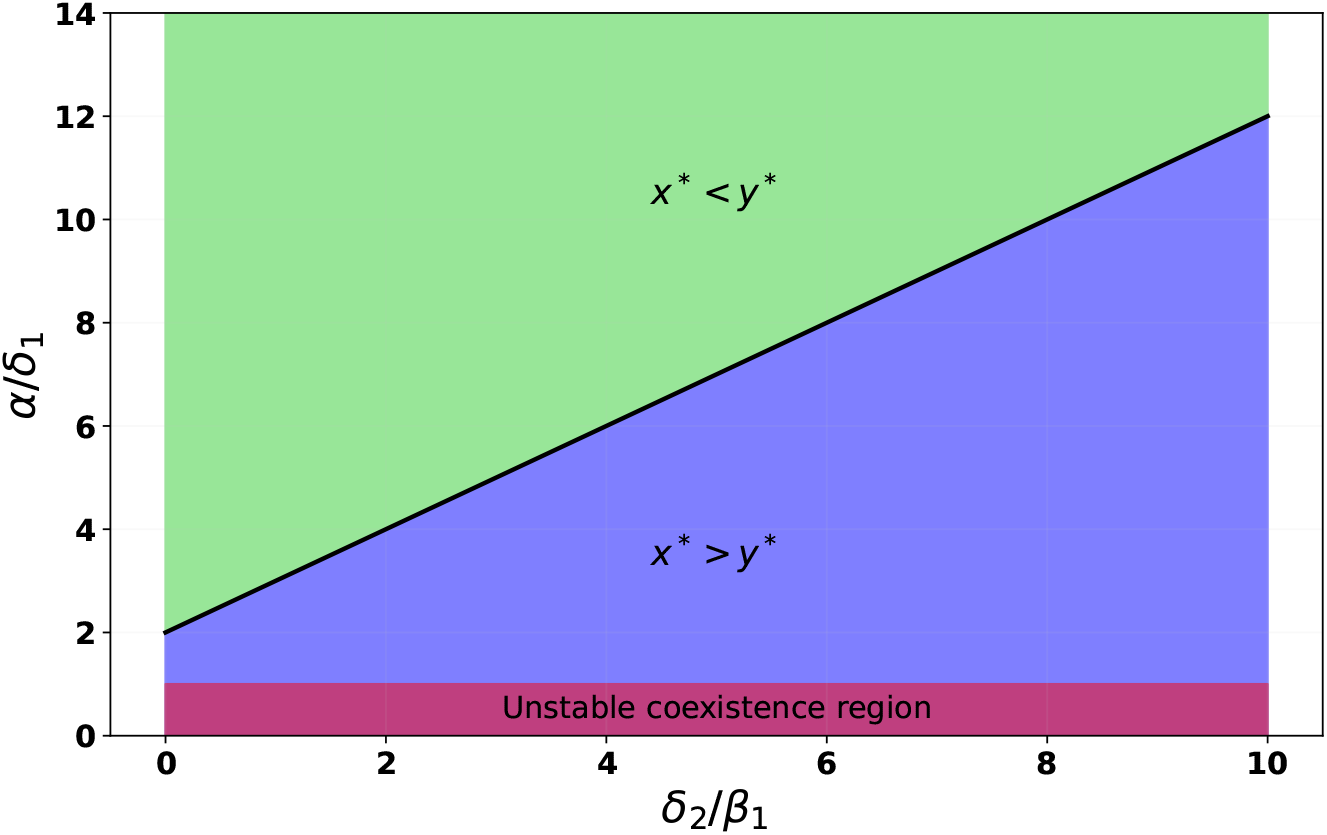
Relative copy numbers of the autonomous-nonautonomous TE interaction. The average number of complexes produced by an autonomous TE, *α/δ*_1_, needs to exceed 1 for stable coexistence to occur (red background). If it exceeds the value 2 + *δ*_2_*/β*_1_ (black line), then nonautonomous TEs are more abundant compared to autonomous TEs (green background), if it is between 1 and 2 + *δ*_2_*/β*_1_ (black line), then autonomous TEs are more abundant (blue background).

## 4 Discussion

We studied a previously described model of the interaction of nonautonomous and autonomous transposable elements (Eq. (5); Xue and Goldenfeld (2016)). To analyze the system, we applied a time scale separation, where the numbers of TEs change on a slow time scale and the intermediate complex, necessary for TE transposition, is comparably short-lived (fast time scale). Based on this time scale separation, we first investigated the stability of the equilibria in the deterministic system and then quantified the stationary distribution of the stochastic model.

In the deterministic model, we find that stable coexistence of the two TEs is possible if the autonomous TE exhibits exponential growth in the absence of the nonautonomous TE (Eq. (10)). In other words, we find that nonautonomous TEs, in form of their transposition parameter *β*_2_, do not directly contribute to the stability of the TE population (see also Appendix D for the more general model with different TE deactivation rates). Yet, it is their presence that keeps the system stable, which is mediated through their dependence on the transposition machinery of autonomous TEs for their mobility. This has prompted their labeling as “hyperparasitic”, “superparasitic” or “parasites of parasites” (Startek et al., 2013; Robillard et al., 2016; Suh, 2019), considering that TEs propagate as parasites of the host. Alternatively, taking the perspective of the TEs instead of the host, one can view the nonautonomous elements as regulators of excessive TE proliferation (Rio, 1991; Hartl et al., 1992), because an increase in their rate of transposition, *β*_2_, results in a reduction of the autonomous TE equilibrium (Eq. (7)). This regulation may serve as a mechanism to prevent bursts of TE proliferation that could have deleterious effects on the host genome (Werren, 2011). We speculate that this controlled proliferation of TEs in a genome may less likely trigger host defenses, potentially leading to long-term persistence of the TE population (Hayward and Gilbert, 2022), though this remains to be studied in models accounting for deleterious fitness effects of TEs and explicitly incorporating host defenses like piRNA clusters (e.g. Kofler, 2019, 2020; Tomar et al., 2023). Thus, from a TE perspective autonomous and nonautonomous elements engage in a form of specialization: autonomous TEs encode the transposition machinery necessary for transposition, while nonautonomous TEs regulate TE proliferation. This specialization supports the ecological perspective of the genome as an ecosystem (Venner et al., 2009; Linquist et al., 2015; Kremer et al., 2020).

Previous population genetic models of autonomous-nonautonomous TE dynamics did not find stable coexistence (Brookfield, 1996; Le Rouzic and Capy, 2006; Startek et al., 2013), but see some exceptions in Le Rouzic and Capy (2006). This contradicts empirical examples, for instance *LINE1* and *Alu* elements in the human genome that seem to be active and coexisting since millions of years in the human genome lineage (Kazazian, 2004). One important difference from our model to previous models is that we account explicitly for the molecular mechanisms of transposition. Our model was motivated by the transposition mechanism of non-LTR retrotransposons. We believe that it can also capture dynamics of LTR retrotransposons, though this remains to be investigated. While including molecular details into the model might already explain the difference to previous results, it is also possible that previous models failed to explain coexistence because in these models autonomous TEs give rise to nonautonomous TEs through mutation. Note, however, that it is possible to find parameter configurations where mutations from autonomous to nonautonomous copies result in stable coexistence, yet it remains unclear how general this result is (Le Rouzic and Capy, 2006). Our assumption of ‘unrelated’ autonomous and nonautonomous TE families is in line with empirical observations, e.g. *LINE1* and *Alu* elements in the human genome and also the RLG *Wilma* and RLG *Sabrina* elements in the wheat genome seem to be unrelated, i.e., they are more than just a couple of mutations away from each other (Kazazian, 2004; Wicker et al., 2021). Another difference to previous models is that we did not model any fitness effects of the TEs on the host. The consequences of this assumption remain to be studied, though we hypothesize that adding weak deleterious fitness effects of TEs on the host will not change the qualitative behavior of our model.

However, it is important to emphasize that not all autonomous-nonautonomous TE interactions will result in long-term coexistence. For example, the empirical study by Robillard et al. (2016) demonstrated autonomous-nonautonomous TE dynamics that did not result in long-term coexistence. They studied the autonomous *Mos1* and nonautonomous *peach* elements of the *mariner* family in *Drosophila melanogaster*. Their findings revealed that when both TEs are present in the same genome, the nonautonomous TEs amplify significantly, indicative of a very large nonautonomous transposition rate *β*_2_ compared to the autonomous transposition rate *β*_1_, which adversely affects the survival and reproductive activity of the autonomous TE. This empirical observation fits with our model prediction because for transposition rates *β*_2_ ≫ *β*_1_, the autonomous equilibrium *x*^*^ = *β*_1_*/β*_2_ converges to zero, which ultimately results in no transposition.

Our main result is the computation of the stationary variance in the stochastic model (Eq. (17)). The stationary distribution, which we derive from a linear noise approximation, is a normal distribution centered around the deterministic equilibrium (Eq. (7)); derivation of the stationary distribution with the more exact central limit theorem gives almost exactly the same result (Eq. (24) in Appendix B). Surprisingly, the covariance matrix in stationarity takes a very specific form in our analysis in the main text (Eq. (16)). We find that the stationary variances of the autonomous and nonautonomous TEs can be rewritten in terms of the covariance between the two TEs and the deterministic equilibria. Importantly, this opens the possibility to test this TE model against data. Given multiple sequences from the same species and a known autonomous-nonautonomous TE pair, e.g. active *LINE1* and *Alu* copy numbers obtained from different human genomic sequences (Hancks and Kazazian, 2012; Dewannieux et al., 2003), one could compare the observed variance of the elements to the predicted form, which is computed from the sample means and the sample covariance.

Additionally, the stationary covariance matrix provides further support to the regulatory role of nonautonomous TEs. Each entry of the matrix scales inversely with the nonautonomous transposition rate *β*_2_. That is, stochastic fluctuations in TE copy numbers are small if *β*_2_ is large. Thus, nonautonomous TEs not only affect the equilibrium value of autonomous TEs but regulate the overall variability of the TE dynamics.

We further investigated the effects of recombination on TE dynamics, particularly on the stationary variance. We find that recombination has no visible impact on TE equilibrium (mean) copy numbers (see Fig. 5) compared to Fig. 3. Moreover, for the stationary variance (Eq. (17)), by comparing the analytical and estimated stationary distributions with the data from the stochastic simulations in the form of histograms, we find that under a low recombination rate (*r* = 0.01), both the analytical and estimated stationary distributions fit well with the data (Fig. 5a). However, as recombination rate increases up to free recombination (Fig. 5b, c), only the estimated stationary distribution fits the data for the autonomous TE. This supports the purpose of the rate-independent covariance matrix (Eq. (17)), i.e., given multiple sequences from the same species and a known autonomous-nonautonomous TE pair, data for the autonomous TE in histogram form can be estimated correctly by computing the TEs mean and covariance. However, for the nonautonomous TE, the fit between both the analytic and estimated stationary distribution and the simulated data progressively diverges. This indicates that as recombination increases, the stationary variance (Eq. (17)), particularly for the nonautonomous TE, does not fit the data. This is explained by recombination reducing covariance (or linkage) between different genomic sites. This implies that our theory is best to be compared to TEs that are predominantly found in lowly recombining regions, which is where TEs tend to accumulate (Dolgin and Charlesworth, 2008; Errbii et al., 2024; Kent et al., 2017). However, it would be desirable to extend our predictions to arbitrary recombination rates to obtain more insight about the exact interplay between transposition and recombination rates.

We also explored the relationship between the copy numbers of the two TE types. We found that it is influenced by the interplay between the average number of complexes (transposition machinery) encoded by the autonomous TE and its transposition rate *β*_1_, as presented in Proposition 1. This interaction explains the mechanisms underlying the varying copy numbers of TEs (Schulman and Wicker, 2013). We discovered that if the transposition rate of the autonomous TE *β*_1_ is low, the average number of complexes required for the nonautonomous TE to be more abundant than the autonomous TE increases. Conversely, if the transposition rate of the autonomous TE is high, the average number of complexes required for the nonautonomous TE to be more abundant is low. These insights help explain why nonautonomous TEs are often more abundant than autonomous TEs (Robillard et al., 2016). Examples include the *Alu* element, which accounts for approximately 1.1 million copies with about 852 active ones, and the *LINE1* element, which accounts for approximately 516 000 copies with about 40 − 50 active ones in the human genome (Hancks and Kazazian, 2012).

Lastly, we also examined the relative variation of TE copy numbers as presented in Proposition 2. We find that when the mean copy number of autonomous TEs is lower than that of nonautonomous TEs, the variance of autonomous TEs is always smaller than that of nonautonomous TEs. Conversely, if the mean copy number of autonomous TEs exceeds that of nonautonomous TEs, no simple prediction of the relationship between stationary variances of the TEs is possible. Variations in TE copy number can also shed light on genome stability, with higher variance indicating greater genomic instability, while lower variance, regulated by nonautonomous TEs or the host, may signify genomic regions with more stable TE content and reduced activity rates (Bhat et al., 2022).

In conclusion, we analyzed a stochastic model of autonomous-nonautonomous TE interactions. This interaction can produce a stable coexistence of both TE types and therefore is a potential mechanism to explain the maintenance of TE transposition in genome lineages over long periods of time. Additionally, our theoretically predicted stationary distribution of TE copy numbers is suited to be compared to empirical data of (active) TE copy numbers because its special form does not require knowledge about TE transposition or deactivation rates.

## Data availability

The Julia and Python codes, data files, and resulting figures are publicly available on GitHub (https://github.com/Ade-omole/TEs_long_term).

## Acknowledgments

We are grateful to the High-Performance Computing service of the University of Münster for providing computational resources on the computer cluster PALMA II.

## Appendix

### A Proof of Proposition 1

*Proof*. Statements (b) and (c) follow directly from statement (a). To prove statement (a), we have

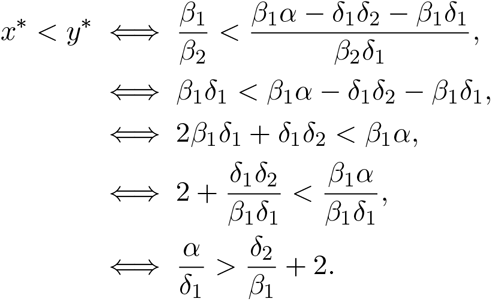

This concludes the proof.

#### Visualization of Proposition 1

In Fig. 6 we show how the average number of complexes produced by an autonomous TE (x-axis) affects the relationship between mean copy numbers of autonomous and nonautonomous TEs.

### B Approximating the stationary distribution with the method from Kang et al. (2014)

Here, we provide a second way to approximate the stationary distribution. Instead of using the equilibrium value *C*^*^ whenever possible, which we did in the main text, we now account for the stochastic nature of the intermediary complex. Under a time scale separation, as we apply it, this method was introduced as a central limit theorem by Kang et al. (2014). We will follow the application of this method as presented in Appendix A of Czuppon and Pfaffelhuber (2018). To this end, we first write down the infinitesimal generator *L*^*V*^ (applied to a auxiliary function *f* (*C, x*^*V*^, *y*^*V*^ )) of the rescaled model as stated in Eq. (2):

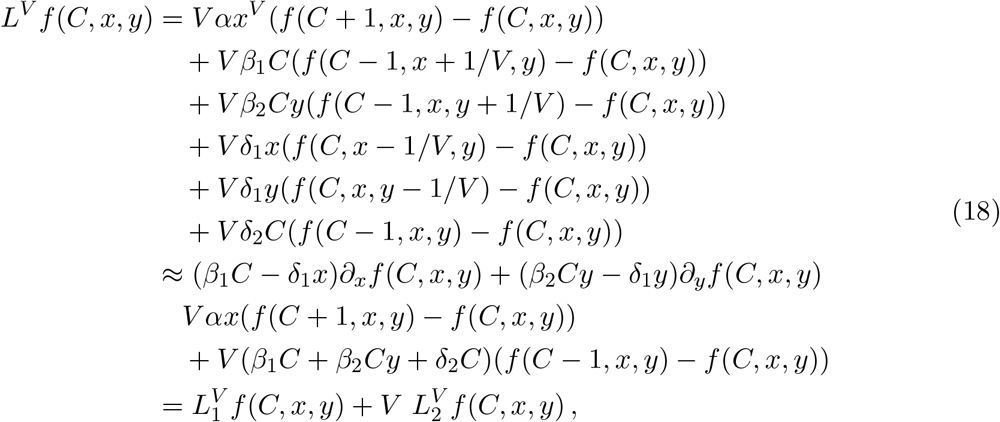

where 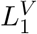 is the infinitesimal generator of the ‘slow’ subsystem and 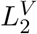 corresponds to the ‘fast’ subsystem. The deterministic dynamics under the time scale separation are given by

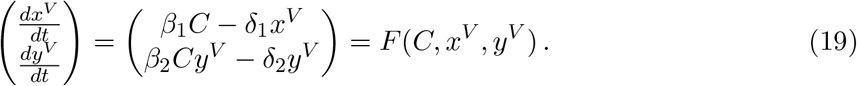

Following step ❷ in Appendix A from Czuppon and Pfaffelhuber (2018), the idea is to find a function *h*(*C, x, y*) so that 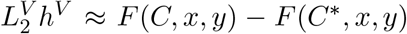. Setting *h*^*V*^ (*C, x, y*) = (−*Cβ*_1_*/*(*β*_1_ + *β*_2_*y* + *δ*_2_), −*Cβ*_2_*y/*(*β*_1_ + *β*_2_*y* + *δ*_2_), we find

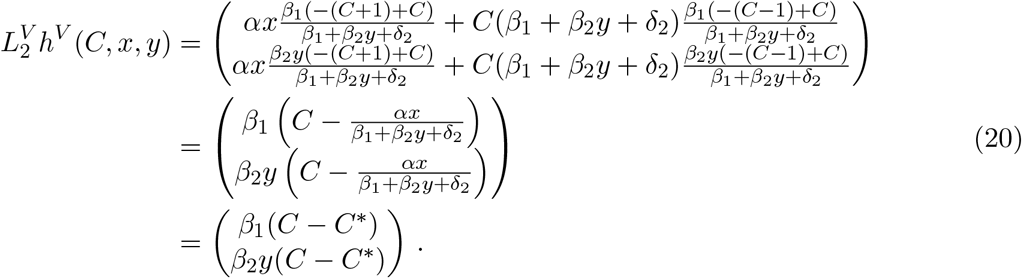

Lastly, we have to compute the quadratic variation (❹ in Czuppon and Pfaffelhuber (2018)). We now denote by 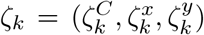 the change in numbers due to the *k*^th^ transition in Eq. (1) and by *λ*_*k*_ the corresponding transition rate. Then, we find that with *z*^⊗2^ = *zz*^⊤^ and 𝒦 the number of transitions, the quadratic variation is given by

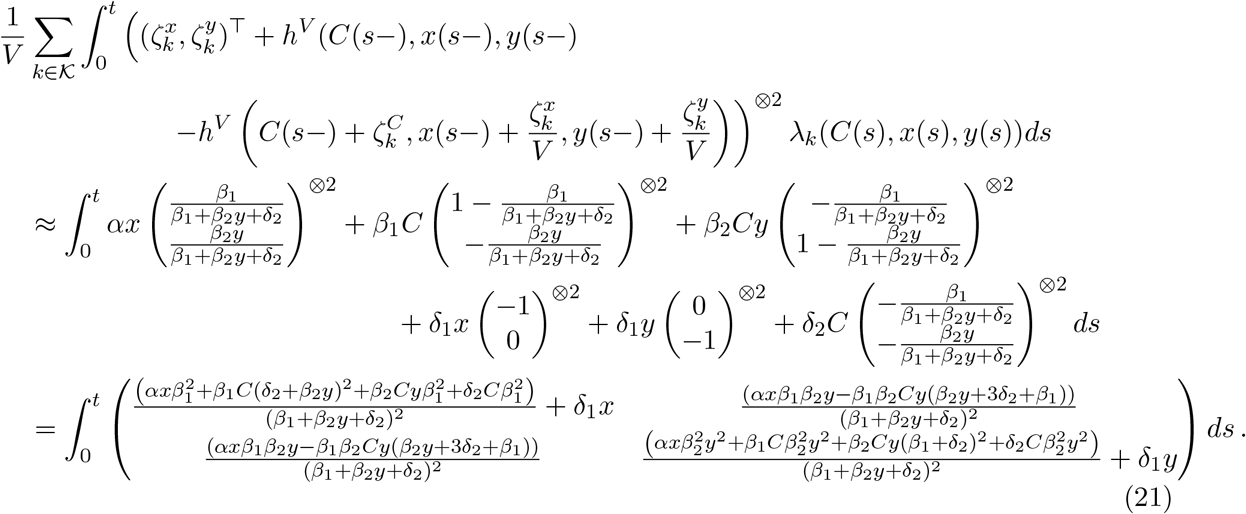

Evaluated in the equilibrium (*C*^*^, *x*^*^, *y*^*^), we obtain for the quadratic variation

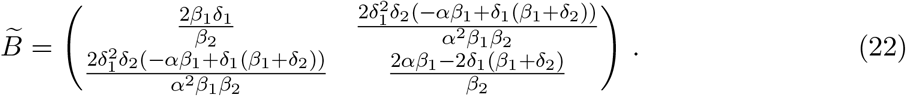

Now following the same steps as in the main text (Eqs. (13) and (14)) with 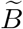 as just defined, we find

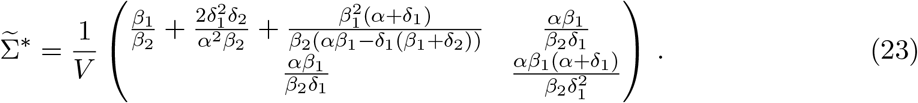

Compared to Eq. (15) only the variance term of the autonomous TE (top left entry) has changed. Again, writing *ξ* = *αβ*_1_*/*(*β*_2_*δ*_1_), we can rewrite the stationary covariance matrix as

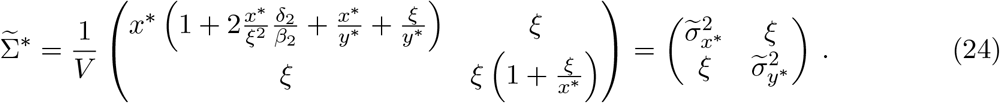

Contrarily to the less accurate method from the main text, we cannot rewrite the stationary variance of the autonomous TE, 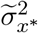, in terms of the steady states and the covariance. Instead, we need knowledge about two parameter ratios, *δ*_2_*/β*_2_ and *α/δ*_1_, if we wanted to compare this stationary variance to data. The difference to the calculated variance in the main text, 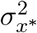 is

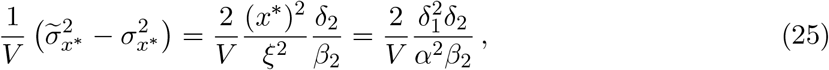

which in our default parameter set amounts to 7.5 × 10^−4^. This term becomes negligible if the rate of TE deactivation, *δ*_1_, is very low, the rate of complex degradation, *δ*_2_, is very low, the rate of complex production, *α*, is very high, or if the rate of nonautonomous TE transposition, *β*_2_, is very high.

### C Proof of Proposition 2

*Proof*. To prove Proposition 2, we first determine the threshold value *x*^*^*/y*^*^ when 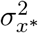 is equal to 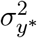. To this end, we simplify

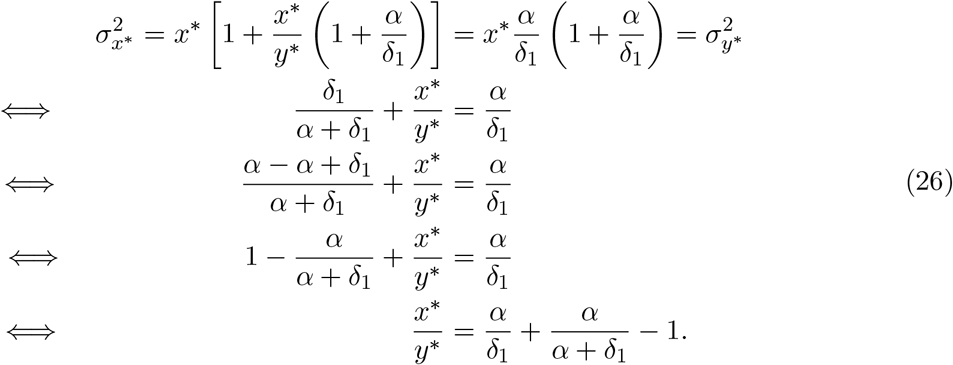

(a) In this case, we have *x*^*^ ≤ *y*^*^. With Proposition 1 (a) and (c), we then also have 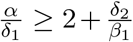. Now, comparing this to Eq. (26), we see that the left hand side is smaller than 1, and the right hand side is larger than 1, so that we find:

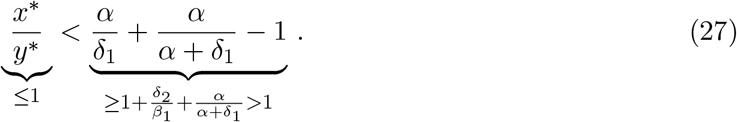

This implies that 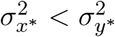.

(b) Here, we have *x*^*^ > *y*^*^. Writing 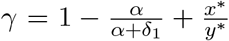, and plugging this into the second last line in Eq. (26), we then directly have that:

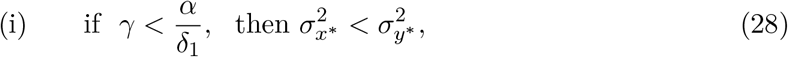

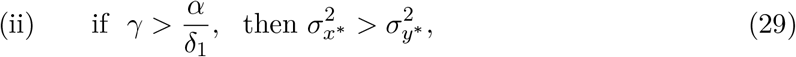

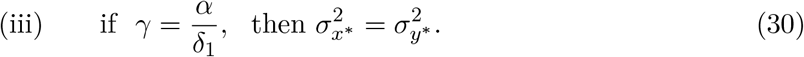

### D Model with different TE deactivation rates

In the main text we have assumed that both TEs are deactivated at the same (scaled) rate *δ*_1_. Here, we study the more general model where we allow for different deactivation rated *δ*_*x*_ and *δ*_*y*_. The model in the main text is recovered by setting *δ*_*x*_ = *δ*_*y*_ = *δ*_1_.

We first note that the equilibrium complex copy number *C*^*^ is unaffected and remains as given in Eq. (3). The stochastic differential equations are then given by

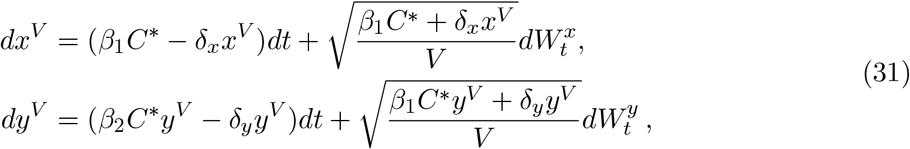

The non-trivial deterministic equilibrium of this model reads

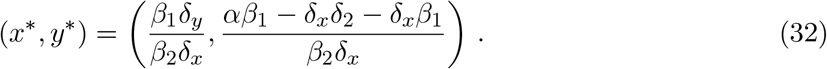

This equilibrium exists if *αβ*_1_ − *δ*_*x*_*δ*_2_ − *δ*_*x*_*β*_1_ > 0, which is the same condition as in Eq. (8) in the main text. Moreover, the equilibrium is locally stable if it exists (determinant-trace condition evaluated in the supplemental *Mathematica* notebook). We can therefore conclude that the TE-specific deactivation rate does not affect the local stability of the system and that the biological interpretations drawn in the main text extend to this more general case.

To derive the stationary distribution, we repeat the same steps as in the main text to find the following stationary covariance matrix:

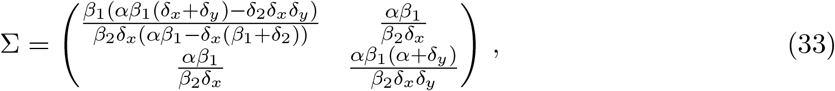

which reduces to Eq. (15) for *δ*_*x*_ = *δ*_*y*_ = *δ*_1_. Again, rewriting the covariance as *ξ* = *αβ*_1_*/*(*β*_2_*δ*_*x*_), we can rewrite the stationary variance of *y* as 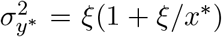. However, we were not able to find a simplification of the stationary variance of *x* as in the main text, which is independent of model parameters. We arrive at

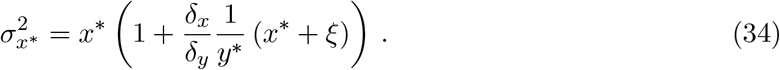

## References

Anderson, D. F. and Kurtz, T. G. Continuous Time Markov Chain Models for Chemical Reaction Networks, page 342. Springer New York, 2011. doi: 10.1007/978-1-4419-6766-41.

Ball, K., Kurtz, T. G., Popovic, L., and Rempala, G. Asymptotic analysis of multiscale approximations to reaction networks. The Annals of Applied Probability, 16(4), 2006. doi: 10.1214/105051606000000420.

Bennett, P. M. Plasmid encoded antibiotic resistance: acquisition and transfer of antibiotic resistance genes in bacteria. British Journal of Pharmacology, 153, 2008. doi: 10.1038/sj.bjp.0707607.

Bennetzen, J. L. and Wang, H. The contributions of transposable elements to the structure, function, and evolution of plant genomes. Annual Review of Plant Biology, 65(1):505–530, 2014. doi: 10.1146/annurev-arplant-050213-035811.

Bhat, A., Ghatage, T., Bhan, S., Lahane, G. P., Dhar, A., Kumar, R., Pandita, R. K., Bhat, K. M., Ramos, K. S., and Pandita, T. K. Role of transposable elements in genome stability: Implications for health and disease. International Journal of Molecular Sciences, 23(14):7802, 2022. doi: 10.3390/ijms23147802.

Boeke, J. D. LINEs and Alus – the polyA connection. Nature Genetics, 16(1):6–7, 1997. doi: 10.1038/ng0597-6.

Brookfield, J. F. Y. Models of the spread of non-autonomous selfish transposable elements when transposition and fitness are coupled. Genetical Research, 67(3):199–209, 1996. doi: 10.1017/s0016672300033681.

Brookfield, J. F. and Badge, R. M. Population genetics models of transposable elements. Genetica, 100(1/3):281–294, 1997. doi: 10.1023/a:1018310418744.

Charlesworth, B. and Langley, C. H. The evolution of self-regulated transposition of transposable elements. Genetics, 112(2):359383, 1986. doi: 10.1093/genetics/112.2.359.

Charlesworth, B. Transposable elements in natural populations with a mixture of selected and neutral insertion sites. Genetical Research, 57(2):127134, 1991. doi: 10.1017/s0016672300029190.

Charlesworth, B. and Charlesworth, D. The population dynamics of transposable elements. Genetical Research, 42(1):1–27, 1983. doi: 10.1017/s0016672300021455.

Cooke, S. L., Shlien, A., Marshall, J., Pipinikas, C. P., Martincorena, I., Tubio, J. M., Li, Y., Menzies, A., Mudie, L., Ramakrishna, M., et al. Processed pseudogenes acquired somatically during cancer development. Nature Communications, 5(1), 2014. doi: 10.1038/ncomms4644.

Cosby, R. L., Chang, N.-C., and Feschotte, C. Host-transposon interactions: conflict, cooperation, and cooption. Genes & Development, 33(1718):1098–1116, 2019. doi: 10.1101/gad.327312.119.

Czuppon, P. and Pfaffelhuber, P. Limits of noise for autoregulated gene expression. Journal of Mathematical Biology, 77(4):1153–1191, 2018. doi: 10.1007/s00285-018-1248-4.

Czuppon, P. and Traulsen, A. Understanding evolutionary and ecological dynamics using a continuum limit. Ecology and Evolution, 11(11):5857–5873, 2021. doi: 10.1002/ece3.7205.

Dewannieux, M., Esnault, C., and Heidmann, T. LINE-mediated retrotransposition of marked Alu sequences. Nature Genetics, 35(1):41–48, 2003. doi: 10.1038/ng1223.

Dolgin, E. S. and Charlesworth, B. The effects of recombination rate on the distribution and abundance of transposable elements. Genetics, 178(4):2169–2177, 2008. doi: 10.1534/genetics.107.082743.

Doolittle, W. F. and Sapienza, C. Selfish genes, the phenotype paradigm and genome evolution. Nature, 284(5757):601–603, 1980. doi: 10.1038/284601a0.

Errbii, M., Gadau, J., Becker, K., Schrader, L., and Oettler, J. Causes and consequences of a complex recombinational landscape in the ant cardiocondyla obscurior. Genome Research, 34(6):863–876, 2024. doi: 10.1098/rstb.2016.0458.

Ethier, S. N. and Kurtz, T. G. Markov Processes: Characterization and Convergence. Wiley, 1986. doi: 10.1002/9780470316658.

Feschotte, C., Swamy, L., and Wessler, S. R. Genome-wide analysis of mariner -like transposable elements in rice reveals complex relationships with Stowaway Miniature Inverted Repeat Transposable Elements (MITEs). Genetics, 163(2):747–758, 2003. doi: 10.1093/genetics/163.2.747.

Gillespie, D. T. Approximate accelerated stochastic simulation of chemically reacting systems. The Journal of Chemical Physics, 115(4):1716–1733, 2001. doi: 10.1063/1.1378322.

Hancks, D. C. and Kazazian, H. H. Active human retrotransposons: variation and disease. Current Opinion in Genetics and Development, 22(3):191–203, 2012. doi: 10.1016/j.gde.2012.02.006.

Hartl, D. L., Lozovskaya, E. R., and Lawrence, J. G. Nonautonomous transposable elements in prokaryotes and eukaryotes. Genetica, 86(13):47–53, 1992. doi: 10.1007/bf00133710.

Hayward, A. and Gilbert, C. Transposable elements. Current Biology, 32(17):R904–R909, 2022.

Kallenberg, O. Foundations of Modern Probability. Springer International Publishing, 2021. ISBN 9783030618711. doi: 10.1007/978-3-030-61871-1.

Kang, H.-W., Kurtz, T. G., and Popovic, L. Central limit theorems and diffusion approximations for multiscale Markov chain models. The Annals of Applied Probability, 24(2), 2014. doi: 10.1214/13-aap934.

Kazazian, H. H. Mobile elements: Drivers of genome evolution. Science, 303(5664):1626–1632, 2004. doi: 10.1126/science.1089670.

Kelleher, E. S., Barbash, D. A., and Blumenstiel, J. P. Taming the turmoil within: New insights on the containment of transposable elements. Trends in Genetics, 36(7):474–489, 2020. doi: 10.1016/j.tig.2020.04.007.

Kent, T. V., Uzunovi, J., and Wright, S. I. Coevolution between transposable elements and recombination. Philosophical Transactions of the Royal Society B: Biological Sciences, 372(1736):20160458, 2017. doi: 10.1098/rstb.2016.0458.

Kofler, R. Dynamics of transposable element invasions with piRNA clusters. Molecular Biology and Evolution, 36(7):1457–1472, 2019. doi: 10.1093/molbev/msz079.

Kofler, R. piRNA clusters need a minimum size to control transposable element invasions. Genome Biology and Evolution, 12(5):736–749, 2020. doi: 10.1093/gbe/evaa064.

Kremer, S. C., Linquist, S., Saylor, B., Elliott, T. A., Gregory, T. R., and Cottenie, K. Transposable element persistence via potential genome-level ecosystem engineering. BMC Genomics, 21(1), 2020. doi: 10.1186/s12864-020-6763-1.

Le Rouzic, A. and Capy, P. Population genetics models of competition between transposable element subfamilies. Genetics, 174(2):785–793, 2006. doi: 10.1534/genetics.105.052241.

Le Rouzic, A., Boutin, T. S., and Capy, P. Long-term evolution of transposable elements. Proceedings of the National Academy of Sciences, 104(49):19375–19380, 2007. doi: 10.1073/pnas.0705238104.

Linquist, S., Cottenie, K., Elliott, T. A., Saylor, B., Kremer, S. C., and Gregory, T. R. Applying ecological models to communities of genetic elements: the case of neutral theory. Molecular Ecology, 24(13):32323242, 2015. doi: 10.1111/mec.13219.

McClintock, B. Controlling elements and the gene. Cold Spring Harbor Symposia on Quantitative Biology, 21:197–216, 1956. doi: 10.1101/sqb.1956.021.01.017.

Montgomery, E. A., Huang, S. M., Langley, C. H., and Judd, B. H. Chromosome rearrangement by ectopic recombination in Drosophila melanogaster : Genome structure and evolution. Genetics, 129(4):1085–1098, 1991. doi: 10.1093/genetics/129.4.1085.

Orgel, L. E. and Crick, F. H. C. Selfish DNA: the ultimate parasite. Nature, 284(5757):604–607, 1980. doi: 10.1038/284604a0.

Otto, S. P. and Day, T. A Biologist’s Guide to Mathematical Modeling in Ecology and Evolution. Princeton Univ. Press, Princeton, NJ, 2007.

Pealba, J. V. and Wolf, J. B. W. From molecules to populations: appreciating and estimating recombination rate variation. Nature Reviews Genetics, 21(8):476492, 2020. doi: 10.1038/s41576-020-0240-1.

Rio, D. C. Regulation of Drosophila P element transposition. Trends in Genetics, 7(9):282–287, 1991. doi: 10.1016/0168-9525(91)90309-e.

Robillard, E., Le Rouzic, A., Zhang, Z., Capy, P., and Hua-Van, A. Experimental evolution reveals hyperparasitic interactions among transposable elements. Proceedings of the National Academy of Sciences, 113(51):14763–14768, 2016. doi: 10.1073/pnas.1524143113.

Schulman, A. H. and Wicker, T. A field guide to transposable elements. Plant Transposons and Genome Dynamics in Evolution, pages 15–40, 2013. doi: 10.1002/9781118500156.ch2.

Startek, M., Le Rouzic, A., Capy, P., Grzebelus, D., and Gambin, A. Genomic parasites or symbionts? Modeling the effects of environmental pressure on transposition activity in asexual populations. Theoretical Population Biology, 90:145–151, 2013. doi: 10.1016/j.tpb.2013.07.004.

Suh, A. Genome size evolution: Small transposons with large consequences. Current Biology, 29(7):R241–R243, 2019. doi: 10.1016/j.cub.2019.02.032.

Suh, D., Choi, E., Yamaziki, T., and Harada, K. Studies on the transposition rates of mobile genetic elements in a natural population of Drosophila melanogaster . Molecular Biology and Evolution, September 1995. doi: 10.1093/oxfordjournals.molbev.a040253.

Tomar, S. S., Hua-Van, A., and Le Rouzic, A. A population genetics theory for piRNA-regulated transposable elements. Theoretical Population Biology, 150:1–13, 2023. doi: 10.1016/j.tpb.2023.02.001.

Tubio, J. M., Li, Y., Ju, Y. S., Martincorena, I., Cooke, S. L., Tojo, M., Gundem, G., Pipinikas, C. P., Zamora, J., Raine, K., et al. Extensive transduction of nonrepetitive DNA mediated by L1 retrotransposition in cancer genomes. Science, 345(6196), 2014. doi: 10.1126/science.1251343.

van Kampen, N. G. Stochastic Processes in Physics and Chemistry. Elsevier, 2007. doi: 10.1016/b978-0-444-52965-7.x5000-4.

Venner, S., Feschotte, C., and Bimont, C. Dynamics of transposable elements: towards a community ecology of the genome. Trends in Genetics, 25(7):317–323, 2009. doi: 10.1016/j.tig.2009.05.003.

Wagstaff, B. J., Hedges, D. J., Derbes, R. S., Campos Sanchez, R., Chiaromonte, F., Makova, K. D., and Roy-Engel, A. M. Rescuing Alu: Recovery of New Inserts Shows LINE-1 Preserves Alu Activity through A-Tail Expansion. PLoS Genetics, 8(8):e1002842, 2012. doi: 10.1371/journal.pgen.1002842.

Wei, W., Gilbert, N., Ooi, S. L., Lawler, J. F., Ostertag, E. M., Kazazian, H. H., Boeke, J. D., and Moran, J. V. Human L1 retrotransposition: cis preference versus trans complementation. Molecular and Cellular Biology, 21(4):1429–1439, 2001. doi: 10.1128/mcb.21.4.1429-1439.2001.

Werren, J. H. Selfish genetic elements, genetic conflict, and evolutionary innovation. Proceedings of the National Academy of Sciences, 108:10863–10870, 2011. doi: 10.1073/pnas.1102343108.

Wicker, T., Stritt, C., Sotiropoulos, A. G., Poretti, M., Pozniak, C., Walkowiak, S., Gundlach, H., and Stein, N. Transposable element populations shed light on the evolutionary history of wheat and the complex coevolution of autonomous and nonautonomous retrotransposons. Advanced Genetics, 3(1), 2021. doi: 10.1002/ggn2.202100022.

Xue, C. and Goldenfeld, N. Stochastic predator-prey dynamics of transposons in the human genome. Physical Review Letters, 117(20), 2016. doi: 10.1103/physrevlett.117.208101.

